# Isolation, spectral characterization and biological activity of fractions of the flavonoid-containing *Gratiola officinalis* L. extract

**DOI:** 10.1101/2020.11.30.404475

**Authors:** Alexander Shirokov, Vyacheslav Grinev, Natalya Polukonova, Roman Verkhovsky, Alena Doroshenko, Dmitry Mudrak, Nikita Navolokin, Anna Polukonova, Alla Bucharskaya, Galina Maslyakova

## Abstract

The extract of *G. officinalis* was separated by five major fractions and the antitumor activity of hydrophobic fractions has been established. For the most active fractions spectral investigations of the main component were performed using UV-Vis electronic spectra, circular dichroism (CD), Fourier-transform infrared spectroscopy (FTIR) and nuclear magnetic resonance (NMR) spectroscopy. On the basis of spectral characteristics, it was demonstrated that the main components of the fractions **4** and **5** of *G. officinalis* providing its biological activity, highly likely, are of flavonoid nature. Based on the NMR data we propose the structure of the aglycon moiety of the main component from fraction **4** as 4,2’,5’-trihydroxychalcone. The fraction **5** is the most active, the antitumor effect of which is implemented through cytotoxic, cytostatic, apoptotic and autophagosomal activity in respect to human kidney cancer tumor cells.

## 1. Introduction

Flavonoids belong to the most promising group of antitumor and preventive herbal remedies, as they have the greatest number of biological effects, including anticancer activity [1]. The organic solvents (ethyl and methyl alcohols of various concentrations, ether, benzene, hexane, chloroform) and purified water are usually used to isolate flavonoids from medicinal raw materials [2]. Thus, the medicinal preparation “Flamin” (Vititekh, Russia), containing the sum of flavonoids, is obtained from flowers of *Helichrysi flos* by extraction with 50% ethanol by the countercurrent method. The extraction is evaporated in a vacuum apparatus to 1/4 of the original volume, the resulting precipitate is dissolved in water. Flavonoids from the aqueous solution are extracted with a mixture of ethyl acetate and ethanol (9 : 1). The extract is dehydrated with dried sodium sulfate and evaporated under vacuum until the solvent is completely removed [3].

*Gratiola officinalis* L., which belongs to the Scrophulariaceae family, also known as Hedge Hyssop and it is found growing wild from Western and Central Europe to Western Asia [4]. *G. officinalis* has diuretic, purgative and vermifuge properties, and as a herbal tea is used to treat chronic gastroenteritis, renal colic, jaundice and intestinal worms [5], however, all parts of the plant are poisonous [6]. The extract of *G. officinalis*, obtained by the author’s method using 95% ethyl alcohol as an extractant (Patent No. 2482863, RF), contains flavonoids, is non-toxic and possesses antitumor, antioxidant, antimicrobial, anti-inflammatory, anti-cahexic and other properties [7–9]. The extract of *G. officinalis* causes the neutralization of free radicals and stabilization of cell membranes [10], affects the immunological status and protects the DNA of cells from the damaging effects of xenobiotics [11], which gives grounds to class it as a modifier of biological reactions that increase the antitumor resistance of the organism as a whole. The chemical composition of *G. officinalis* extract is not fully understood. It is known that the composition of raw materials *G. officinalis* includes gratiogenin, 16-hydroxygratiogenin, cucurbitacins-E, glycosides gratiogenin-3 beta-D-glucoside, gratioside, elaterinide, lignans, traces of alkaloids, coumarin and mannitol derivatives, as well as betulinic acid [12–17].

Analysis of the *G. officinalis* extract by using gas chromatography-mass spectrometry (GC-MS) showed the presence of the bioflavonoid quercetin, as well as residual content of volatile compounds: 4-vinyl-2-methoxyphenol, 2,3-dihydro-3,5-dihydroxy-6-methyl-4*H*-pyran-4-one, 2,3-dihydrobenzofuran, 3-furancarboxylic acid, 5-hydroxymethyl-2-furaldehyde, 4-propylphenol, pyrocatechol, ethyl ester of benzoylacetic acid, homovanillic acid, benzoic acid and gallic acid [18].

The authenticity of *G. officinalis* extract is confirmed by qualitative reactions to the presence of flavonoids and the absence of alkaloids. To determine the presence of flavonoids, the Synod sample was used, reactions with aluminum chloride, with alkali, with sodium hydrogen carbonate, with iron-ammonium alum and reactions to the presence of alkaloids were carried out: with Wagner-Bouchard reagent, with 1% pycrynic acid solution, with phosphorus-molybdenum acid solution, with silicon-ulframic acid [19].

Further studies of the chemical composition of the active fractions and the mechanisms of the antitumor action of *G. officinalis* extract are necessary for its promotion as a new medicinal anticancer agent.

The aim of this work was to separate the extract of *G. officinalis* into individual fractions and identify the most biologically active fractions, and to study their chemical composition and biological activity on the culture of human kidney cancer A498 cells.

## 2. Results and discussion

### 2.1. The separation of extract of G. officinalis into fractions

As far the extract of *G. officinalis* was preliminarily treated by nonpolar solvent, at the initial step of our work, we have used semi-preparative HPLC method to improve the quality of the separation of the whole extract. The dry mass of 1 mL of the extract used was 107.7 mg, and the total mass of the dried fractions obtained by semi-preparative HPLC was 57.7 mg. The masses of individual fractions are shown in Table S1 and typical chromatograms are given on Figures S1-S5. To evaluate the relative content of the components of the redissolved the extract in the eluent we used gravimetric method. The mass of dry residue, obtained for more detailed fractionation from 1 mL of the extract solved in the eluent, was determined to be 0.1032 g.

The chromatogram profile of the whole extract of *G. officinalis* in the used gradient elution mode mostly contain peaks with retention times assigned to the most hydrophobic compounds (Fig. 1). Fractions collecting time ranges were corrected to provide the presence of the most abundant components in single fractions, in particular, after 16 min.

**Figure 1.**
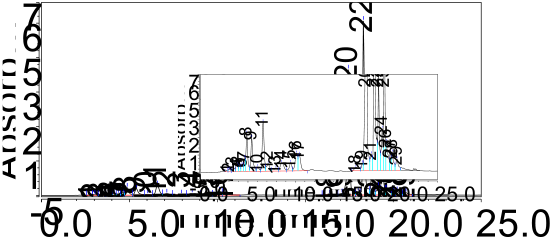
The normalized to highest peak chromatogram profile of the whole extract of *G. officinalis* (in the insert is the same chromatogram showing intensities of less intensive peaks).

All fractions were analyzed on the analytical C18 column using the same elution method. We used semi-preparative and analytical columns with the same sorbent and manufactured by one vendor for better scalability. The fractions containing one major and significantly less other components were chosen for further investigation. Fractions with the same composition were combined.

As a result, the fractions were given new designations and their dry masses were combined in Table S4. Finally, *G. officinalis* extract was divided into five separate fractions (signed as **1-5**) for the further investigations.

### 2.2. Spectral analysis of fractions 4 and 5 of the extract of G. officinalis

The most hydrophobic fractions **4** and **5** of *G. officinalis* extract revealed high biological activity and were additionally characterized by a set of spectroscopic data (UV, CD, FTIR and NMR spectroscopy) to establish the probable structure of the components.

#### 2.2.1. UV-Vis spectral analysis

The use of diode array detector (DAD) in HPLC allows us to obtain UV-Vis spectra of the components of the fractions to make an additional validation and assignment of polyphenolic compounds, which have characteristic electron spectra profiles. For polyphenolic compounds of flavonoid class, rings A and B are considered as two main chromophores, conjugated or not conjugated depending on the unsaturation degree in ring C. Generally, in electronic spectra of flavonoids, two main absorption bands are observed, which intensities ratio and positions of the maxima change widely depending on the presence, location and nature of the substituents. Often, these two bands are called I and II (chromophores of ring B and A, respectively [20]). The UV-Vis spectra of the major components of fractions **4** and **5** in the range of 200-400 nm in acetonitrile-water mixture are shown in (Fig. 2).

**Figure 2.**
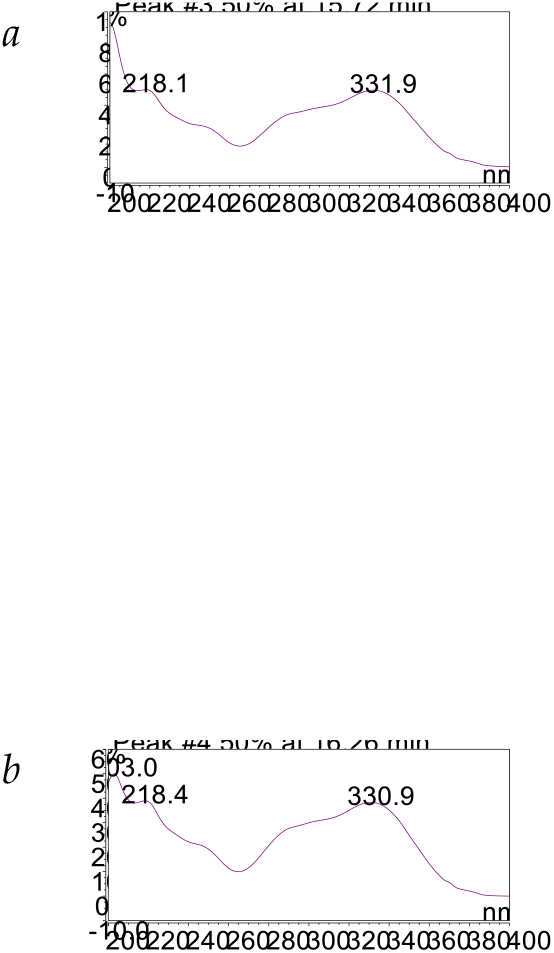
UV-spectrum of fractions **4** (*a*) and **5** (*b*) of the *G. officinalis* extract.

As can be seen, spectra of main components in both fractions are almost identical slightly differing by the position of maximum near 200 nm. In the spectra, there are three more pronounced bands centered at 200-203, 218 and 331-332 nm and two less pronounced bands with maxima at approximately 250 and 290 nm. It is reasonable to assign peaks at 331-332 nm in fractions **4** и **5** to band I of putative flavonoid indicating its flavone nature. The complex structure of the spectra caused probably by overlapping of the conjugated fragments of molecules close to the chemical structure of the extract is detected. The profile of the spectra is highly similar to that of flavonoids, which have two substituted aromatic rings in their structure, one of which is fused with pyran ring (Fig. 3).

**Figure 3.**
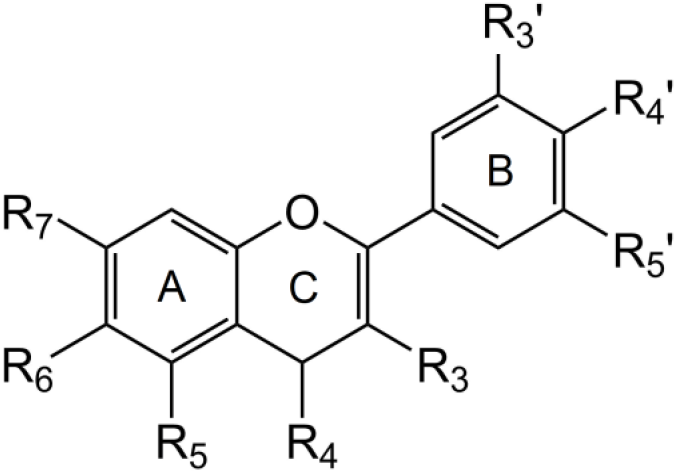
General structure of flavonoids showing the designation of A, C and B rings.

Due to the retention time match, previously, we have made an assignment for this component to quercetin. A comparison of the UV-Vis spectra profiles of quercetin standard sample and the unknown component of the extract showed their difference (data not shown).

#### 2.2.2. Circular dichroism (CD) analysis

It is well known that optical active isomers, especially enantiomers, may act in different way demonstrating non-equivalent biological activity. With the aim to evaluate the presence of the optical active compounds in most hydrophobic fractions of the extract we used circular dichroism (CD) spectroscopy. The most common optical active substituent in flavonoids is carbohydrate moiety, which have no chromophores itself. In contrast, an aglycon moiety is a clearly pronounced chromophore which allows one to register CD spectra (Fig. 4).

**Figure 4.**
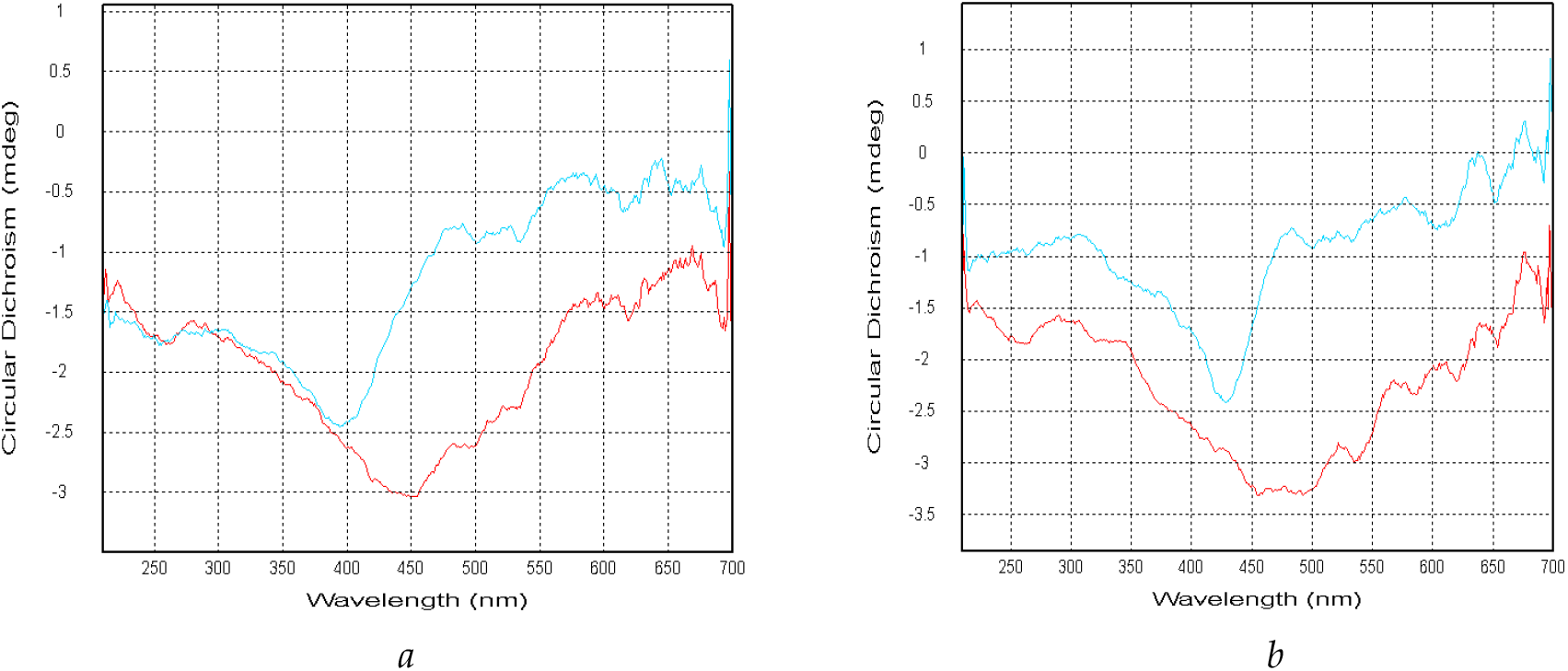
Circular dichroism (CD) spectrum of fractions **4** (A) and **5** (B). Optical path length is 2 mm (red line) and 10 mm (blue line).

Fractions **4** and **5** demonstrate weakly pronounced optical activity. In the CD spectra of both fractions there is a negative Cotton effect near 400-430 nm when measured in the cell with an optical path length of 10 mm and red shifting about 50 nm when measured in the cell with an optical path length of 2 mm. On the one hand, the CD spectra confirm that in the structure of the optical active components of the fractions near the chromophore there is an asymmetry due to the presence of probably chiral carbon atom. On the other hand, in the 200-400 nm range which is characteristic for flavonoid-like molecules chromophore, in CD spectra there is no significant Cotton effect which allows us to conclude that the optical activity of the fractions is not due to the asymmetry in flavonoids molecules.

#### 2.2.3. FTIR analysis

Analysis of the functional groups of the components of the *G. officinalis* extract was performed using FTIR spectroscopy. The analysis of FTIR spectra of the selected *G. officinalis* fractions (Fig. 5) allows us to speak about the presence in the fraction **5** of components containing hydrogen-bonded hydroxyl groups. The stretching vibrations assigned to OH groups at 3419 cm^−1^ is significantly red-shifted by about 20 cm^−1^ in comparison to the similar absorption band in fraction **4**, which is registered at 3439 cm^−1^. Absorption bands assigned to aliphatic C-H groups in the range of 2857-2983 cm^−1^ for both fractions are practically of the same profile. In the absorption region of carbonyl groups in the spectra of both fractions there is a superposition of several bands that differ in relative intensity. Thus, in fraction **4** there is an intensive absorption at 1633 cm^−1^ with a shoulder at 1686 cm^−1^, while in fraction **5** the most intensive absorption is observed at 1682 cm^−1^, and overlapping bands in this area do not reveal apparent maxima.

**Figure 5.**
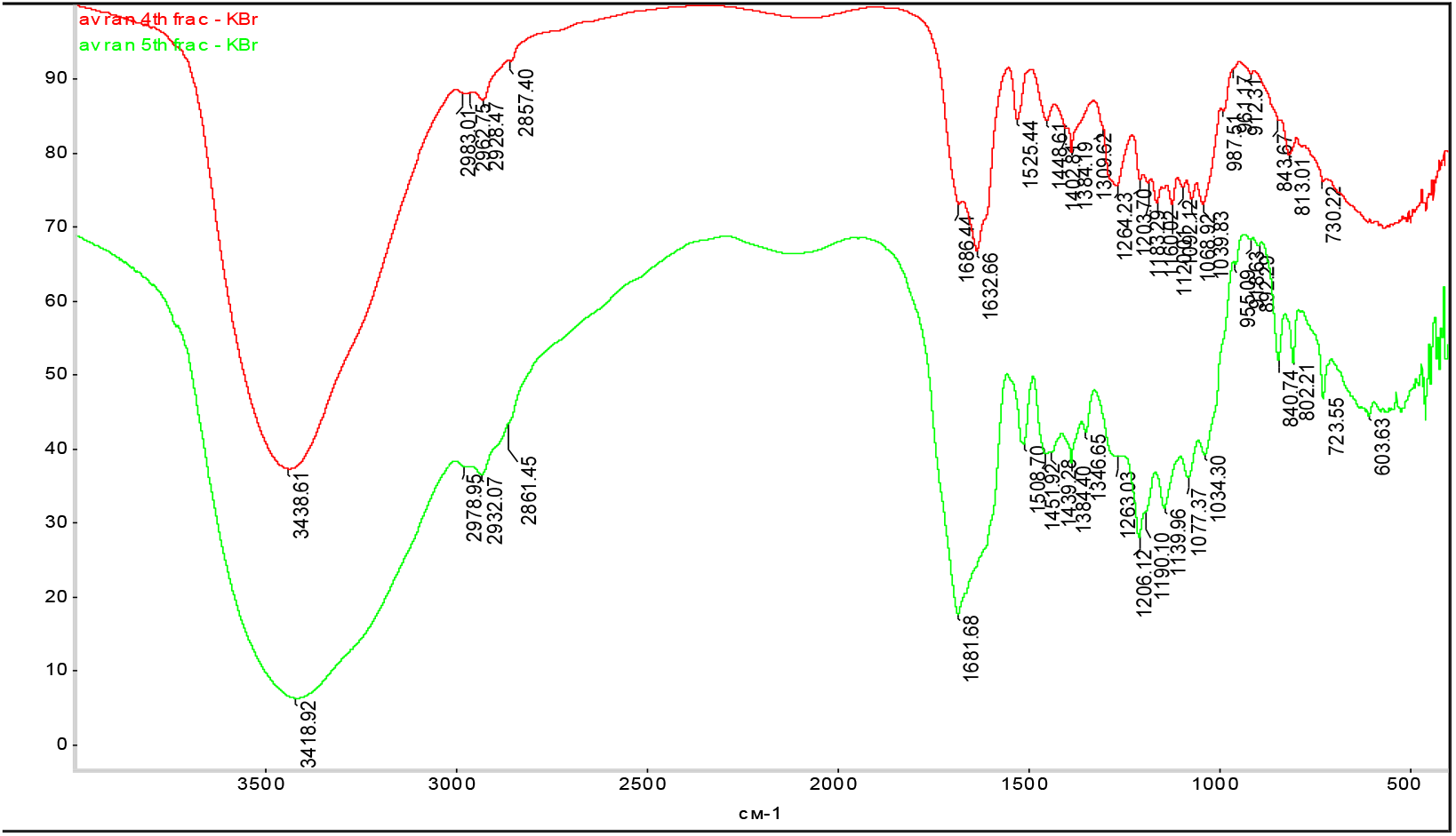
The FTIR spectra of *G. officinalis* extract fractions **4** (red line) and **5** (green line).

Thus, the fractions contain components in whose molecules there are carbonyl groups of different chemical nature. In the fingerprints region there are absorption bands typical for aromatic rings of different degrees of substitution which is in harmony with the structures of flavonoids.

#### 2.2.4. Nuclear magnetic resonance (NMR) analysis

To go further in the analysis of functional groups for fraction **4** (Fig. 6) it was possible to register NMR spectra. In the 6-8 ppm region signals of aromatic protons are observed. One can notice the characteristic doublets of doublets revealing cross-peaks in the COSY spectrum, confirming that they belong to different spin systems of benzene rings. Such structural fragments are typical for different flavonoids. In addition, in 4.2-3.2 ppm region there are well-resolved multiplets that may be assigned to aliphatic protons associated with electron-acceptor substituents. However, it is known that proton signals from carbohydrates are also often found in this region, which requires further analysis, since the retention time of the major component in fraction **4** is typical for non-glycosylated flavonoids. At the same time, some examples are known, when glycosylation of flavonoid for a certain hydroxyl group insignificantly affected the overall polarity of the molecule, which was expressed in a small change in chromatographic characteristics in comparison with the aglyconic form (for example, mono- and diglycosides of naringenin).

**Figure 6.**
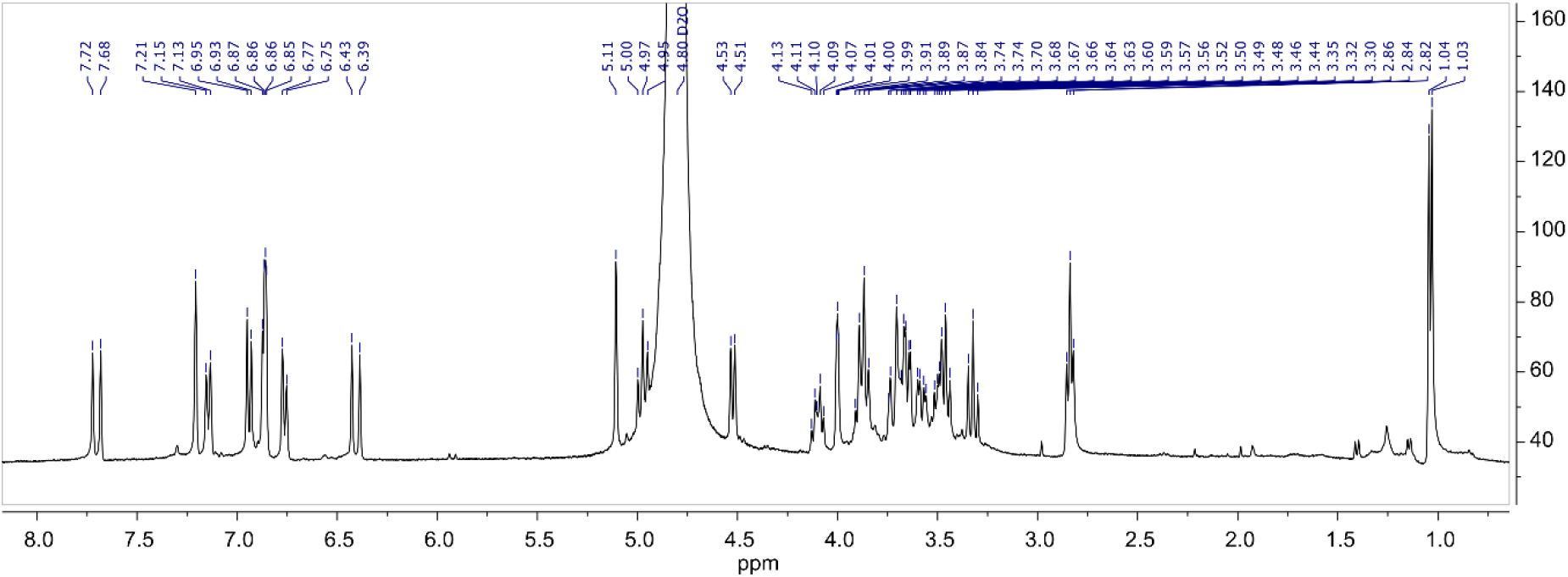
The ^1^H NMR spectrum of the *G. officinalis* extract fraction **4**.

Two doublets at 7.70 and 6.41 ppm both with *J* = 15.9 Hz revealing in the ^1^H-^1^H COSY spectrum (Fig. 7*a*) corresponding cross-peaks belong to protons of AX system at the double bond of chalcone moiety [21]. Two doublets at 7.14 and 6.94 ppm both with *J* = 8.3 Hz were assigned to protons of the *para*-substituted B-ring of the molecule. The presence in the ^1^H-^1^H COSY spectrum of corresponding cross-peaks confirms their proximity. Another ABX system appears as two doublets at 6.76 with *J* = 8.2 Hz and 6.86 ppm with *J* = 8.3 Hz. The last signal is interfering with the doublet at 6.86 ppm with *J* = 2.3 Hz which was also confirmed by the presence of their cross-peaks the ^1^H-^1^H COSY spectrum. Slightly broaded singlet at 7.21 ppm reveals cross-peaks at 6.94/7.21 in the ^1^H-^1^H TOCSY spectrum (Fig. 7*b*) as well as at 7.21/115.3 ppm in the ^1^H-^13^C HSQC spectrum and shows no correlation in the ^1^H-^1^H COSY spectum. The chemical shift value of the bounded carbon along with the position of this signal in the ^1^H the spectrum allows us to assign this signal to aromatic system of the molecule. On the other hand, the relative down-fielded chemical shift value of 7.21 ppm allows us to give preference to 1,2,5-substitution pattern in the A ring instead of 1,2,4-substitution due to the more probable position of the signal of the proton between two phenolic hydroxyl groups in relative up-field region, near 6.20 ppm (for example, for 4,2’,4’-trihydroxychalcone this signal appears at 6.35 ppm [22]).

**Figure 7.**
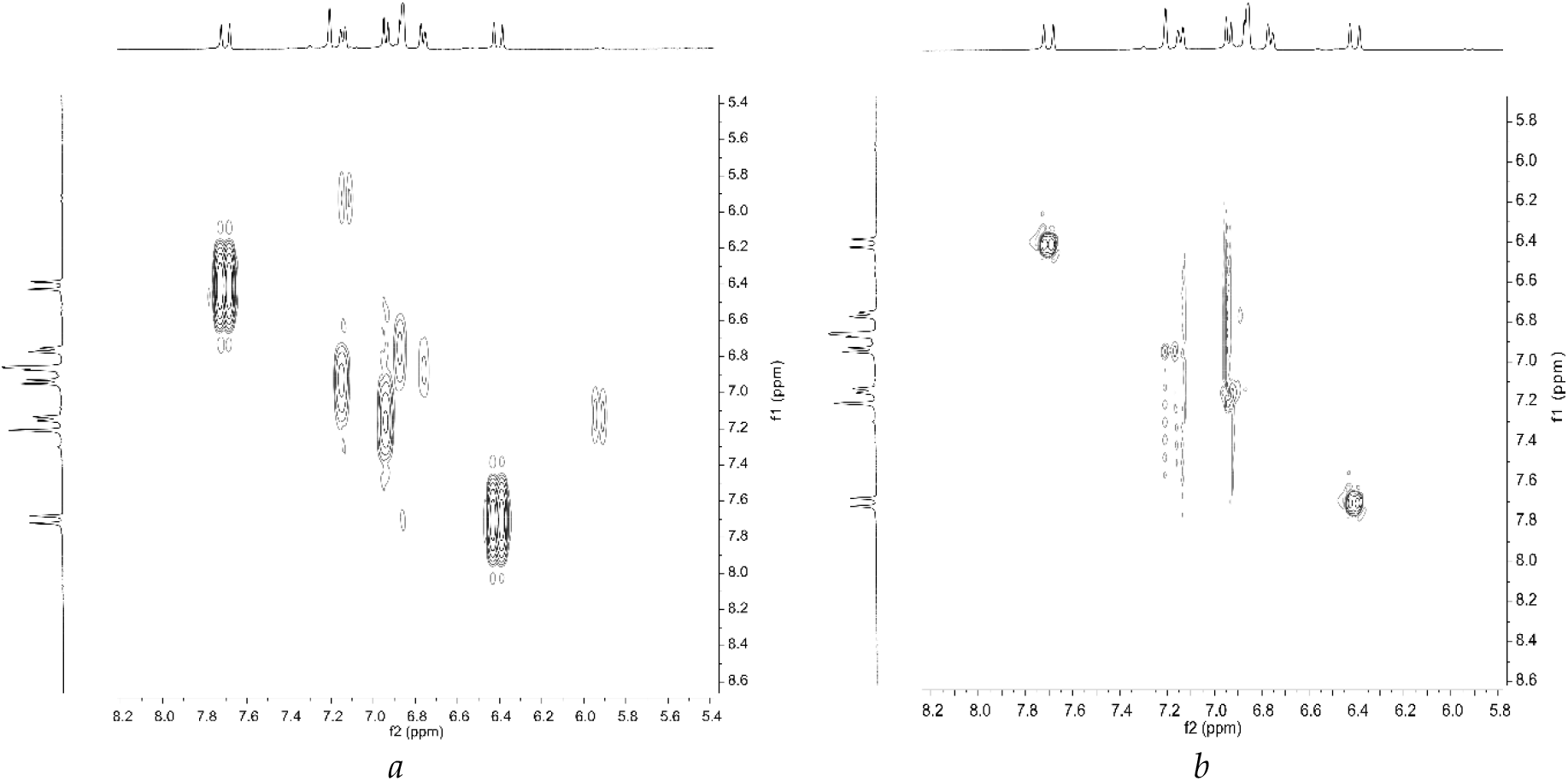
The ^1^H-^1^H COSY (*a*) and TOCSY (*b*) spectra of the *G. officinalis* extract fraction **4**. Diagonal signals are suppressed for clarity.

Methylation of the protons of the phenolic hydroxyl groups is a common modification of the flavonoid molecules, so, we have attempted to find their signals in the NMR spectrum. In the up-field region there are no characteristic singlets at around 3.8 ppm assigned to the methoxy group protons that confirms the absence of the methoxyl substituents in the molecule of the putative flavonoid. Due to the deuterium exchange we can’t watch the signals from labile protons of the phenolic hydroxyl groups.

According to the set of ^1^H and 2D NMR spectra we can conclude that the core moiety of the main component of fraction **4** belongs to class of chalcones and can be matched as 1-(2,5-dihydroxyphenyl)-3-(4-hydroxyphenyl)prop-2-en-1-one (or 4,2’,5’-trihydroxychalcone) with probable substitution of the phenolic hydroxyl groups (Fig. 8). The NMR spectra of 4,2’,5’-trihydroxychalcone collected in different solvents were described recently [22–24]. As it can be seen, in the ^1^H NMR spectrum collected in acetone-*d*6 the signal of the proton at C-6 in the A ring of 4,2’,5’-trihydroxychalcone appears as a doublet with *meta* coupling constant of 3.0 Hz which in case of D_2_O spectrum collapsed into a broaded singlet. This assignment is also in harmony with FTIR spectra which contain bands centered at around 3400 and 1630 cm^−1^ that confirms the presence of an O–H···O=C system [21]. On the other hand, this molecule contains no stereocenters, so the observable optical activity should be assigned with the probable hydrocarbon substituent.

**Figure 8.**
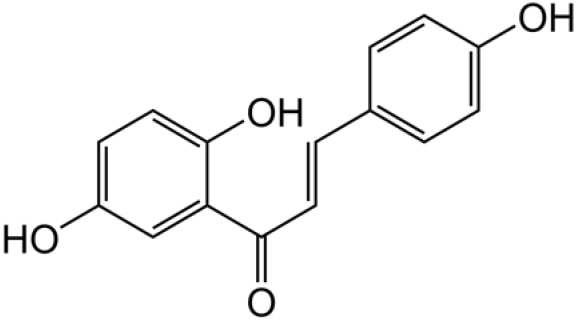
Putative structure of the aglycon moiety of the main component of the *G. officinalis* extract fraction **4**.

### 2.3. Comparison of biological activity of five fractions of G. officinalis extract according to cytomorphological indices on human kidney cancer cells A498

At the next step of our work, we have performed an evaluation of antitumor activity of five collected fractions on human kidney cancer cells to reveal the most active and prominent one.

#### 2.3.1. Fraction 1 and Fraction 3

Values of cytomorphological indices of human kidney cancer cells A498 did not differ from values of these indices in control, both in 24 hours and 48 hours (Table S5, Fig. S11). The data obtained by us testify to the absence of cytostatic and cytotoxic activity of fraction **1** and **3**.

#### 2.3.2. Fraction 2

Under the action of fraction **2**, in comparison with the control, the difference was revealed only in the total number of cells in the field of view and only after 48 hours, namely a decrease in the total number of cells in the field of view was observed ((Table S5, Fig. S11). The obtained data indicate the presence of weak cytostatic activity of fraction **2** with respect to tumor cells of A498 line.

#### 2.3.3. Fraction 4

Under the action of fraction **4**, in comparison with the control, there were revealed differences only in the indicators indicating a cytostatic effect, namely, a decrease in the total number of cells in the field of view and the number of living cells and only after 48 h (Table S5). The data obtained indicate the presence of cytostatic activity of fraction **4** in respect to tumor cells of line A498.

#### 2.3.4. Fraction 5

The most pronounced biological activity was established for fraction **5**.

##### Cytotoxic activity

The number of dead cells and the ratio of the number of dead cells to the total number of cells in the field of view after 24 hours are 10 and 19 times lower respectively (Fig. 9, S6, Table S5), which indicates the presence of cytotoxic activity of this fraction in relation to tumor cells.

**Figure 9.**
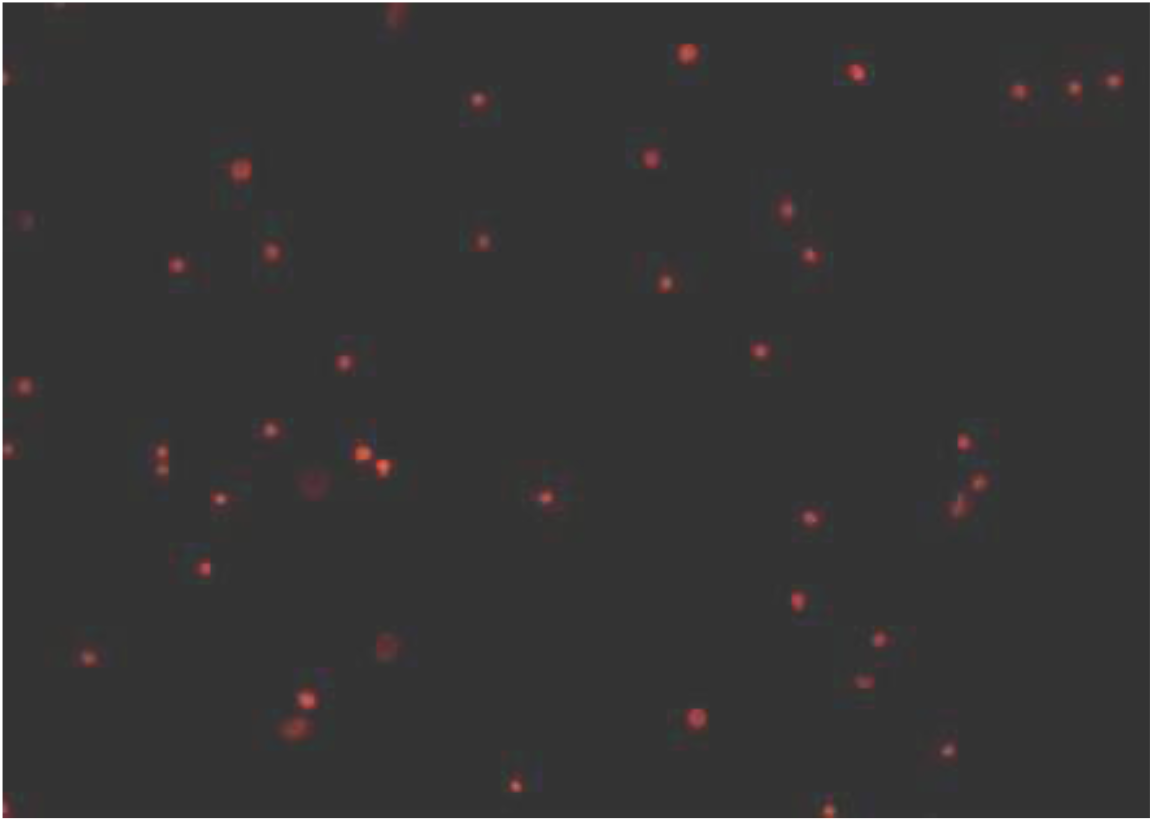
Cytotoxic effect of fraction 5 of G. officinalis extract in human kidney cancer cells A498 Colored with iodine propidium, dead cells are colored red.

##### Cytostatic activity

Under the action of fraction **5**, there were revealed maximum differences of cell indices in comparison with the control, testifying to the cytostatic action of fraction **5** in respect to tumor cells of A498 line. We observed the reduction of total number of cells in the field of view - number of living cells, both in 24 hours and 48 hours, and also the reduction of ratio of living cells to total number of cells in the field of view only in 24 hours (Table S5). At the same time, the reduction of the total number of cells in the field of view under the influence of fraction **5**, increases from 1.6 times in 24 h to 4.1 times to 48 h, as compared with the control; and the reduction of the ratio of living cells to the total number of cells in the field of view was from 3.7 to 10 times to 48 h (Table S5).

Thus, two fractions of *G. officinalis* extract - fraction **4** and fraction **5** - have a pronounced biological activity. Fraction **4** is characterized by its cytostatic action. Fraction **5** is characterized by both cytotoxic and cytostatic action on human tumor cells.

## 3. Materials and Methods

### 3.1. Material

*G. officinalis* extract was obtained according to the original author’s method (Polukonova et al., Patent for invention RUS 2482863 02/15/2012). The investigated composition of biologically active substances (BAS) from *G. officinalis* is one stripped off extract, yellow-brown color, mixed with water and ethyl alcohol in any ratio.

The authenticity of *G. officinalis* extract is confirmed by qualitative reactions for the presence of flavonoids and the absence of alkaloids. We used a sample of the Synod, reactions with aluminum chloride, with alkali, with sodium bicarbonate, with iron-ammonium alum and reactions to the presence of alkaloids: with Wagner-Bouchard reagent, with 1% picricric acid solution, with phosphorus-molybdenum acid solution, with silicon-ulframic acid [19].

### 3.2. Preliminary semi-preparative HPLC fractionation of the extract

Several mL of a mixture consisted of 15% acetonitrile and 85% water with the addition of trifluoroacetic acid (TFA, Sigma-Aldrich, USA) to pH 2.5 to suppress dissociation of labile protons were added to the dried extract, 1 mL was selected for fractionation, 1 mL - for determining the total mass of the dry extract.

At semi-preparative separation ten subfractions were collected manually by time, each of them was analyzed on the analytical C18 HPLC. Chromatograms of all 10 subfractions are presented on Figures S1-S10. Identical subfractions were combined to give five major fractions according to Table S2. All collected fractions were concentrated under reduced pressure on the rotary evaporator Heidolph to a minimum volume and lyophilized using Virtis benchtop dryer. The masses of dry fractions of extracts are given in Table 3.

One mL of selected extract was centrifuged, supernatant was used for fractionation. The fractions were collected manually by time, every 5 min. 5 fraction collection passes were performed on Dionex Ultimate 3000 HPLC (Thermo Scientific, USA) using Nucleodur HTec C18 VP 125/10 column (Macherey-Nagel, Germany) with average particle diameter of 5 μm. Injection volume was 100 μL and eluent flow rate was 2 mL/min. The volume of each collected fraction was 10 mL.

### 3.3. Preparation of samples for extraction of ten fractions of G. officinalis extract

Dried extract was completely dissolved in 10 ml of a mixture of 1:1 acetonitrile and aqueous TFA solution with pH 2.5. The solution of 1 mL was selected to determine the mass of contained dissolved substances.

### 3.4. Analytical HPLC separation

The extract was analyzed using reverse phase HPLC on a Dionex Ultimate 3000 chromatograph (Thermo Scientific, USA) using a Nucleodur HTec C18 VP 125/10 semi-preparative column (Macherey-Nagel, Germany), with an average particle diameter of 5 μm and pores 100 Å and a geometry of 150 × 3.0 mm. Flow rate was 2 mL/min, total analysis time was 25 min.

Chromatographic extract was taken under conditions of gradient elution (Solvent A - acetonitrile of HPLC grade (Panreac, Spain), Solvent B - trifluoroacetic acid (TFA, Sigma Aldrich, USA) solution (pH 2.5): the composition of the mobile phase was changed as follows: 0-10 min. - 15% A, 85% B; 10-19 min. - 15→70% A, 85→30% B; 19-20 min. - 70% A, 30% B; 20-22 min. - 70→15% A, 30→85% B; 22-25 min. - 15% A, 85% B. Total flow was 2 mL/min. Injection volume of the sample was 50 μL. Detection was performed at wavelengths of 342, 360, 320 nm. Chromatograph control and data analysis were performed by Chromeleon version 7.1.2.1478 (“Thermo Scientific”, “Dionex”, USA).

The collected fractions (50 mL each) were vaporized to the minimum volume on the rotary evaporator at a water bath temperature of 42 °С. For the analysis of chromatographic profiles of the obtained fractions, 20 μL from each fraction were selected, the volume was brought to 500 μL by solution of 15% acetonitrile and 85% water (with addition of TFA to pH 2.5). The analysis was performed on the chromatograph using Macherey-Nagel Nucleodur HTec C18 analytic column, average particle diameter was 5 μm with pores 100 Å, geometry of 150 × 3.0 mm. The remaining fractions were lyophilized.

### 3.5. UV-Vis analysis

The electronic spectra of components of the extract were obtained using Dionex™ UltiMate™ DAD 3000 1024-element diode-array detector (Thermo Scientific, USA) in the 200-400 nm range.

### 3.6. FTIR analysis

FTIR spectra were recorded on a Nicolet 6700 Fourier spectrometer (Thermo Scientific) equipped with an absorber of carbon dioxide and water vapors. Samples were grinded with KBr (FTIR grade, Sigma-Aldrich, USA) and pressed at reduced pressure into pellets. Spectra were collected in the 4000-400 cm^−1^ range with spectral resolution of 4 cm^−1^. At least 64 scans were summarized every single spectrum.

### 3.7. Circular dichroism analysis

Circular dichroism (CD) spectra of aqueous solutions of fractions 4 and 5 were recorded on a Chirascan spectrometer (Applied Photophysics, Great Britain) equipped with a Peltier thermostatable cuvette compartment, at 20 °C in a quartz cuvette 115F-QS 10 × 2 mm (Hellma Analytics, Germany), using two optical paths - 2 and 10 mm.

### 3.8. NMR analysis

The ^1^H (400 MHz) and 2D NMR spectra were registered on the Varian 400 spectrometer [Agilent Technologies (formerly, Varian), USA] in D_2_O, the internal standard was sodium trimethylsilylpropanesulfonate (DSS).

### 3.9. Cell lines

The culture of human kidney carcinoma A 498 tumor cells was obtained from the Bank of Cell Lines of National Medical Research Center of Oncology n.a. N.N. Blokhin" of the Russian Ministry of Health. A498 line cells were grown in DMEM medium with addition of 10% serum of embryonic calf (HyClone, USA), penicillin-streptomycin (PanEco, Russia), L-glutamine (PanEco, Russia), sodium pyruvate, vitamins and amino acids (PanEco, Russia) at 37°C and 5% CO2. The cells were removed from culture bottles with Versen solution. Cells at 70% - 80% of monolayer were used for experiments.

*Cytotoxic activity* was assessed by the following indicators: number of dead cells and ratio of dead cells to total number of cells. The activity was observed when the values of at least one of these indicators were reduced. *Cytostatic activity* was evaluated by the following indicators: total number of cells in field of view, number of living cells, ratio of living cells to total number of cells, number of dividing cells, ratio of dividing cells to number of living cells. Activity was observed when the values of at least one of these indicators decreased.

### 3.10. Statistical analysis

All statistical analysis is performed using Microsoft Office Excel software. Normal distribution of indicators in groups was checked using Shapiro-Wilk criterion. The distribution did not correspond to the normal one, therefore the non-parametric Cruckele-Wallace criterion was used. Reliability was established at *p* < 0.005.

## 4. Conclusions

Thus, the fractions in the antitumor extract of *G. officinalis* have been established, providing its activity – fraction **4** and fraction **5** with probably flavonoid nature. The fraction **5** is the most active, the antitumor effect of which is implemented through cytotoxic, cytostatic, apoptotic and autophagosomal activity in respect to human kidney cancer tumor cells.

The analysis of spectroscopic and chromatographic data for fractions **4** and **5** allows us to state that their main component of flavonoid nature is very likely, and at this stage of the study can not exclude the presence of carbohydrate components, which may explain, in particular, the presence of some optical activity in the spectra of CD, as well as the presence of signals in the area characteristic of carbohydrates, among others. Based on the NMR data we can propose that the structure of the aglycon moiety of the main component from fraction **4** should be assigned as 4,2’,5’-trihydroxychalcone. The collected data set may require additional methods to confirm the structure of individual components of the *G. officinalis* fraction, which is responsible for their biological effect.

## Supplementary Materials

The following are available online at www.mdpi.com/xxx/s1, Figure S1: title, Table S1: title, Video S1: title.

## Author Contributions

Conceptualization, P.P. and G.M.; methodology, V.G.; software, D.M.; validation, V.G. and N.N.; formal analysis, A.S.; investigation, V.G., R.V. and A.P.; resources, A.S.; data curation, A.B.; writing—original draft preparation, V.G. and N.P.; writing—review and editing, A.S.; visualization, R.V.; supervision, G.M.; project administration, A.S.; funding acquisition, N.P. All authors have read and agreed to the published version of the manuscript.

## Acknowledgments

HPLC analysis and fractionation were performed at the Center for Collective Use “Symbiosis” of the Federal State Institution of Science of the Institute of Biochemistry and Physiology of Plants and Microorganisms of the Russian Academy of Sciences (IBPPM RAS) within the research theme No. AAAA-A19-119011890162-3. Cell culture investigations performed at the Laboratory of Cell Technologies of Center for Collective Use of Saratov State Medical University n.a. V.I. Razumovsky.

## Sample Availability

Sample of the extract of *G. officinalis* is available from the authors.

## References

1. Chahar, M.K.; Sharma, N., Dobhal, M.P.; Joshi, Y.C. Flavonoids: A versatile source of anticancer drugs Pharmacogn Rev. 2011, 5(9), pp. 1–12.

2. Georgievskii, V.P.; Komissarenko, N.F.; Dmitruk, S.E. Biologically active substances of medicinal plants [in Russian], 3rd ed.; Nauka: Novosibirsk, Russia, 1990; 333 p.

3. Technology of dosage forms [in Russian], Ed. Ivanova, L., Medicina: Moscow, 1991; 544 p.

4. Bown, D. The Royal Horticultural Society New Encyclopedia of Herbs & Their Uses. Dorling Kindersley: London, 2002; 488 p.

5. Launert, E. The Hamlyn guide to edible and medicinal plants of Britain and Northern Europe. Hamlyn: London, 1989; 288 p.

6. Bown, D. Encyclopaedia of Herbs and their Uses. Dorling Kindersley: London, 1995; 424 p.

7. Navolokin, N.A.; Polukonova, N.V.; Maslyakova, G.N.; Bucharskaya, A.B.; Durnova, N.A. Effect of extracts of *G. officinalis* and Zea mays on the tumor and the morphology of the internal organs of rats with transplanted liver cancer. Russian Open Medical Journal. 2012, 1(2), pp.1–4.

8. Navolokin, N.A.; Mudrak, D.A.; Bucharskaya, A.B.; Matveeva, O.V.; Tychina, S.A.; Polukonova, N.V.; Maslyakova, G.N. Effect of flavonoid-containing extracts on the growth of transplanted sarcoma 45, peripheral blood and bone marrow condition after oral and intramuscular administration in rats. Russian Open Medical Journal. 2017, 6(3), p.304.

9. Polukonova, N.V; Navolokin, N.A; Raĭkova, S.V.; Masliakova, G.N.; Bucharskaia, A.B.; Durnova, N.A.; Shub, G.M. Anti-inflammatory, antipyretic and antimicrobial activity of flavonoid-containing extract of *G. officinalis* L. Eksp Klin Farmakol. 2015, 78(1), pp. 34–38.

10. Navolokin, N.A.; Ivlichev, A.V.; Mudrak, D.A.; Afanas’eva, G.A.; Polukonova, N.V.; Tychina, S.A.; Bucharskaya, A.B.; Maslyakova G.N. Influence of flavonoid-containing extract (*G. officinalis* l.) on the content of vitamin E and intensity of peroxidation processes in the blood of rats with transplanted liver cancer PC-1. Eksp Klin Farmakol. 2017, 80(10), pp. 40–43.

11. Kurchatova, M.N.; Durnova, N.A.; Polukonova, N.V. The effect of flavonoids extracts on the dioxydin induction of micronuclei in red blood cells of outbred white mice. [in Russian] Proceedings of Voronezh State University. Series: Chemistry. Biology. Pharmacy. 2014, 2, pp.58–65.

12. Rudolf, T; Annemarie, H. Triterpene, Über Triterpene, II. Mitteil.: Gratiosid, ein Triterpenglykosid aus *G. officinalis* L. in: Berichte der deutschen chemischen Gesellschaft Weinheim, Wiley-VCH, 1952; 85 (11), pp. 1067–1077.

13. Borodin, L.I.; Litvinenko, V.I.; Kurinnaya, N.V. New flavonoid C-glycosides from *G. officinalis*. Chem. Nat. Compd. 1970, 6(1), pp. 19–24.

14. Rothenburger, J.; Haslinger, E. Caffeic acid glycoside esters from *G. officinalis* L. Eur. J. Org. Chem. 1994, 11, pp.1113–1115.

15. Kaya, G.I.; Melzig, M.F. Quantitative determination of cucurbitacin E and cucurbitacin I in homeopathic mother tincture of *G. officinalis* L. by HPLC. Pharm. Die. 2008, 63(12), pp. 851–853

16. Zia-Ul-Haq, M.; Kausar, A.; Shahid S.A.; Qayum, M.; Ahmad, S.; Khan, I. Phytopharmacological profile of *G. officinalis* Linn.: A review. Journal of Medicinal Plants Research. 2012, 6(16), pp. 3087–3092.

17. Ali, L.; Rizvi, T.S.; Ahmad, M.; Shaheen, F. New iridoid glycoside from *G. officinalis* (Scrophulariaceae). Journal of Asian Natural Product Research, 2012, 14(12), pp.1191–1195.

18. Polukonova, N.V.; Durnova, N.A.; Kurchatova, M.N.; Navolokin, N.A.; Golikov, A.G. Chemical analysis of the new biological active composition from medicative herb hedge-hissop (*G. officinalis* L.) [in Russian] Chemistry of plant raw material, 2014, 4, pp. 165–173.

19. Komarova, M.N.; Nikolaeva, L.A.; Regir, V.G.; Teslov, L.S.; Kharitonova, N.P.; Chatokhina, R.K. Phytochemical analysis of medicinal plant raw materials. Method. instructions for laboratory classes [in Russian], Ed. by K.F. Blinova, Sankt-Peterburg. 1998. 60 p.

20. Korolkin, D.Yu.; Abilov, J.A.; Muzychkina, R.A.; Tolstikov, G.A. Natural flavonoids. [in Russian]Novosibirsk: Academic Publishing House “Theo”, 2007. 232 p.

21. Panichpol, K.; Waterman, P.G. Novel Flavonoids From The Stem Of *Popowia Cauliflora*. Phytochemistry, 1978, 17, pp. 1363–1367.

22. Deodhar, M.; Black, D.StC.; Kumar, N. Acid catalyzed stereoselective rearrangement and dimerization of flavenes: synthesis of dependensin. Tetrahedron, 2007, 63, pp. 5227–5235.

23. Lim, S.S.; Jung, S.H.; Ji, J.; Shin, K.H.; Keum, S.R. Synthesis of flavonoids and their effects on aldose reductase and sorbitol accumulation in streptozotocininduced diabetic rat tissues. Journal of Pharmacy and Pharmacology, 2001, 53, pp. 653–668.

24. Moorthy, N.S.H.N.; Singh, R.J.; Singh, H.P.; Gupta, S.D. Synthesis, Biological Evaluation and *In Silico* Metabolic and Toxicity Prediction of Some Flavanone Derivatives. Chem. Pharm. Bull., 2006, 54(10), pp. 1384–1390.

