## Supplemental Table 1 for "Isolation, spectral characterization and biological activity of fractions of the flavonoid-containing *Gratiola officinalis* L. extract"

**Table S1.** Masses of individual fractions obtained by previous fractioning

| Fraction, time, min | Weght, mg |
| --- | --- |
| 0-5 | 37,5 |
| 5-10 | 3,9 |
| 10-15 | 2,7 |
| 15-20 | 12,4 |
| 20-25 | 1,2 |

**Table S2.** Time interval at previous semi-preparrative separation of the fractions

| № | 1. | 2. | 3. | 4. | 5. | 6. | 7. | 8. | 9. | 10. |
| --- | --- | --- | --- | --- | --- | --- | --- | --- | --- | --- |
| Time interval | 2.50-5.50 | 5.50-6.30 | 6.30-7.30 | 7.30-8.80 | 8.80-14.00 | 15.00-16.00 | 16.00-16.30 | 16.30-16.55 | 16.65-17.65 | 17.65-20.00 |

**Table S3.** Previously obtained fractions of G. officinalis extract

| № previously obtained fractions | m (empty eppendorf), g | m(eppendorf+dry substance), g | m(dry substance), mg |
| --- | --- | --- | --- |
| 1 | 0.8902 | 0.9395 | 49.3 |
| 2 | 0.9012 | 0.9020 | 0.8 |
| 2(2) | 0.8696 | 0.8702 | 0.6 |
| 3 | 0.8887 | 0.8893 | 0.6 |
| 4 | 0.8756 | 0.8766 | 1.0 |
| 4(2) | 0.9089 | 0.9096 | 0.7 |
| 5 | 0.8889 | 0.9004 | 11.5 |
| 6 | 0.9180 | 0.9278 | 9.8 |
| 7 | 0.8698 | 0.8778 | 8.0 |
| 7(2) | 0.9009 | 0.9047 | 3.8 |
| 8 | 0.8905 | 0.8927 | 2.2 |
| 9 | 0.8881 | 0.8885 | 0.4 |
| 9(2) | 0.8891 | 0.8919 | 2.8 |
| 10 | 0.9378 | 0.9420 | 4.2 |
| 10(2) | 0.8758 | 0.8778 | 2.0 |

**Figure S1.** Chromatogram of fraction 1 of G. officinalis extract

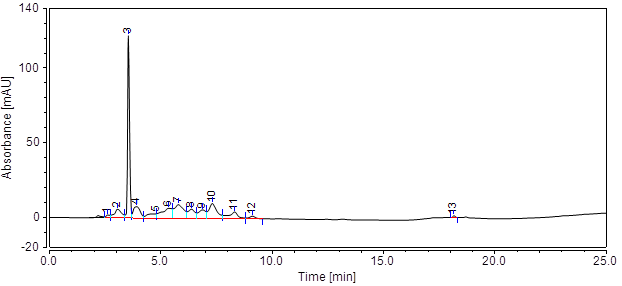

**Figure S2.** Chromatogram of fraction 2 of G. officinalis extract

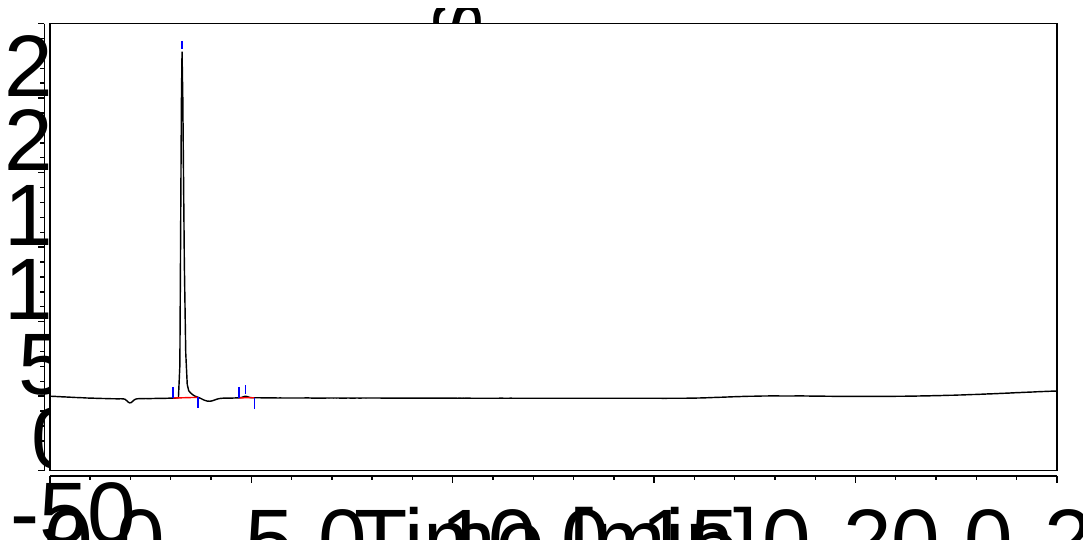

**Figure S3.** Chromatogram of fraction 3 of G. officinalis extract
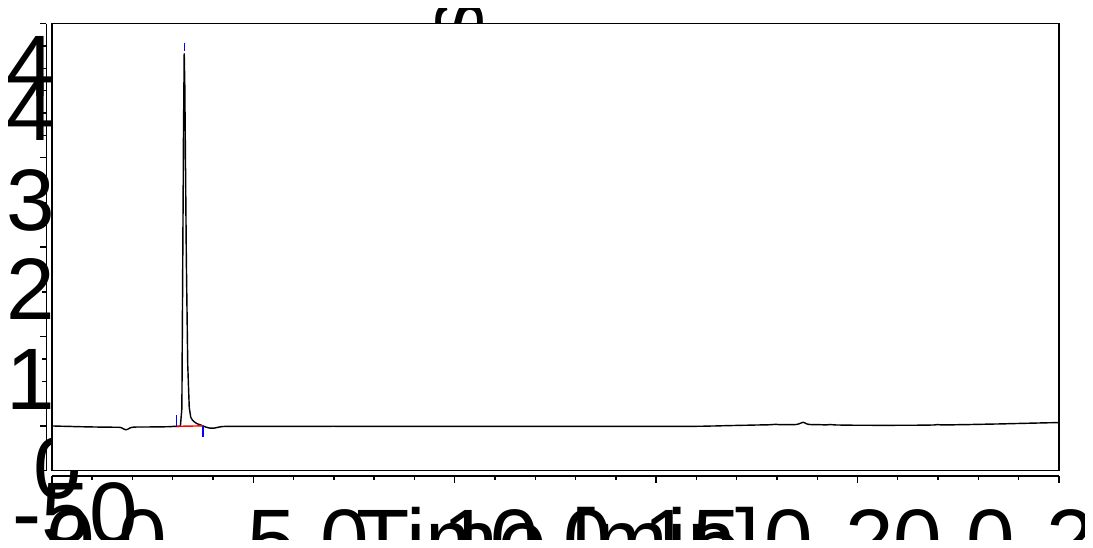

**Figure S4.** Chromatogram of fraction 4 of G. officinalis extract

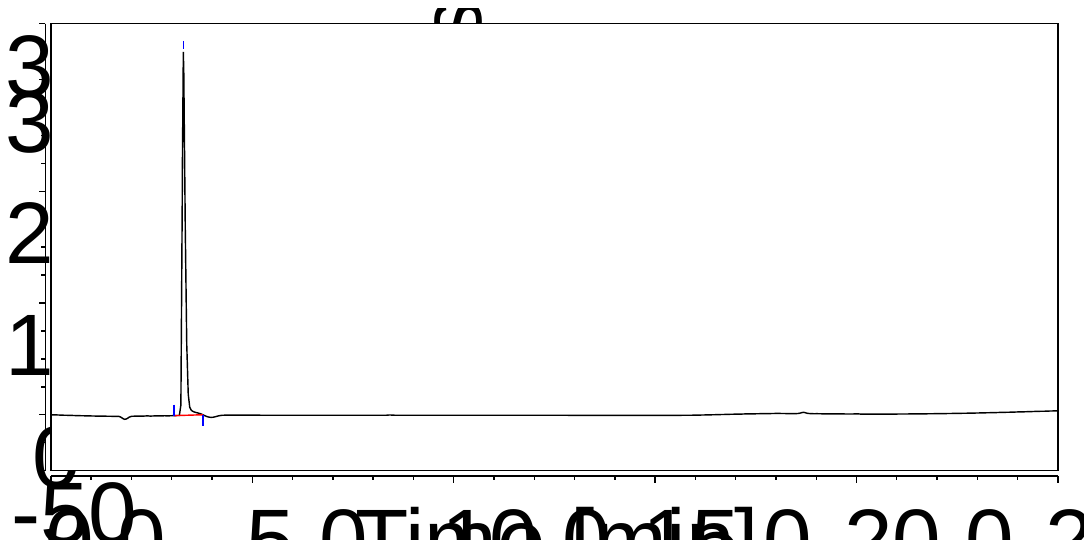

**Figure S5.** Chromatogram of fraction 5 of G. officinalis extract

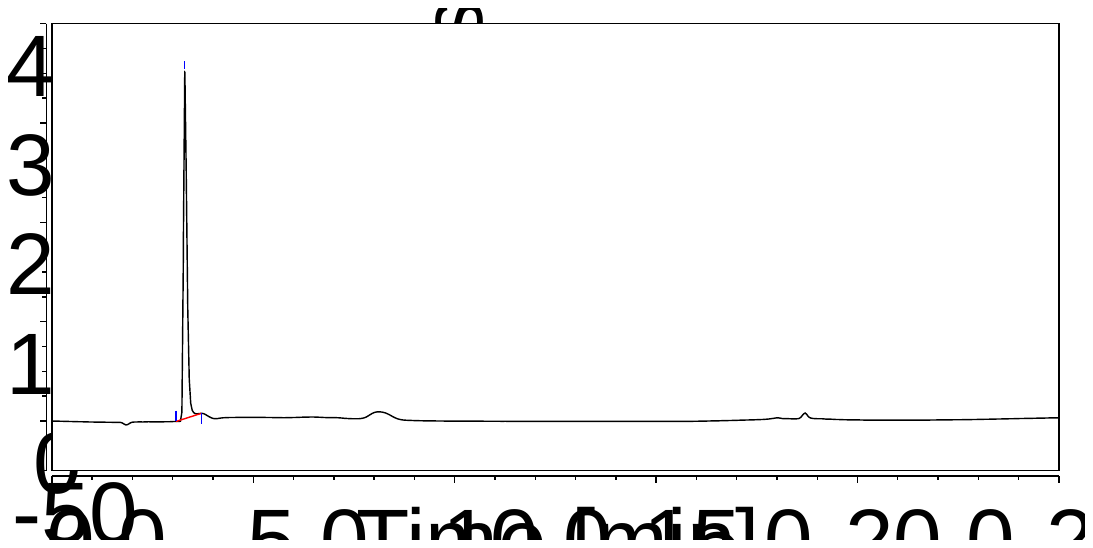

**Figure S6.** Chromatogram of fraction 6 of G. officinalis extract

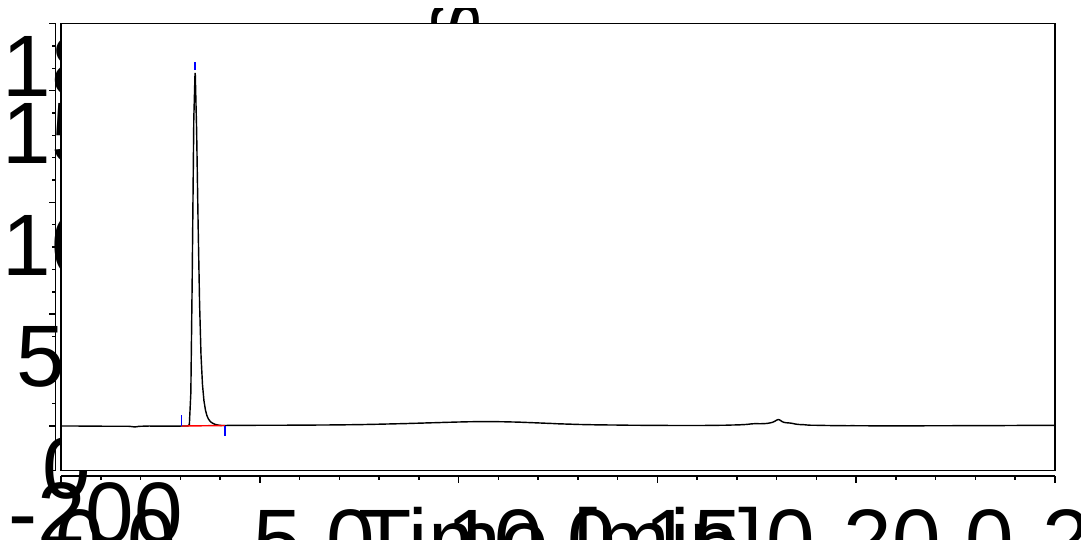

**Figure S7.** Chromatogram of fraction 7 of G. officinalis extract

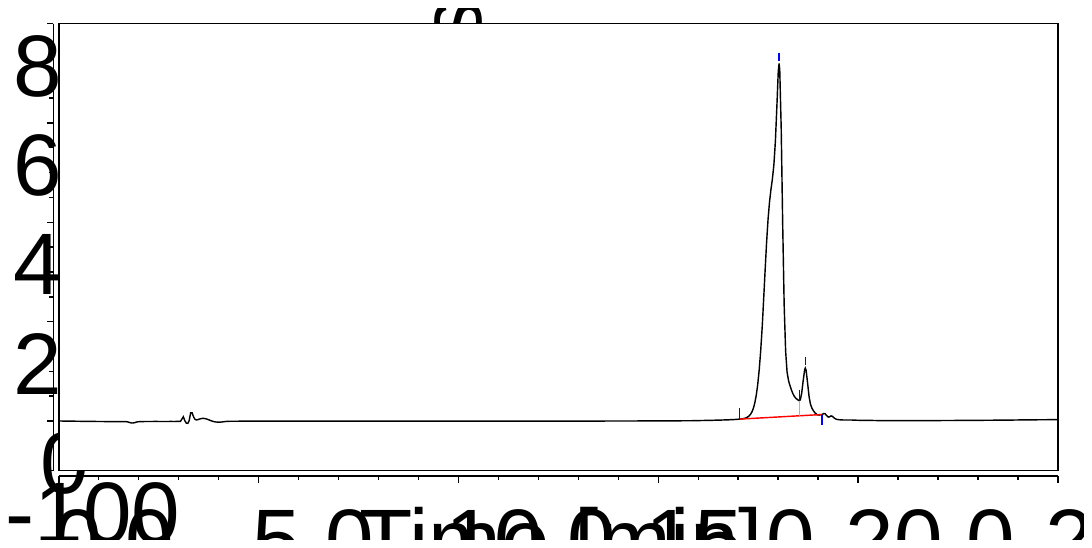

**Figure S8.** Chromatogram of fraction 8 of G. officinalis extract

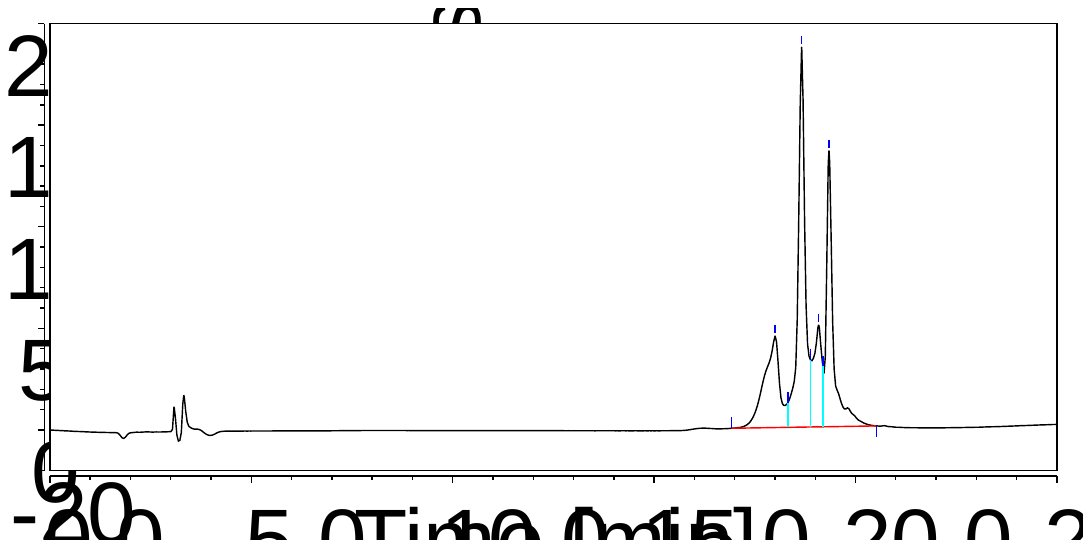

**Figure S9.** Chromatogram of fraction 9 of G. officinalis extract

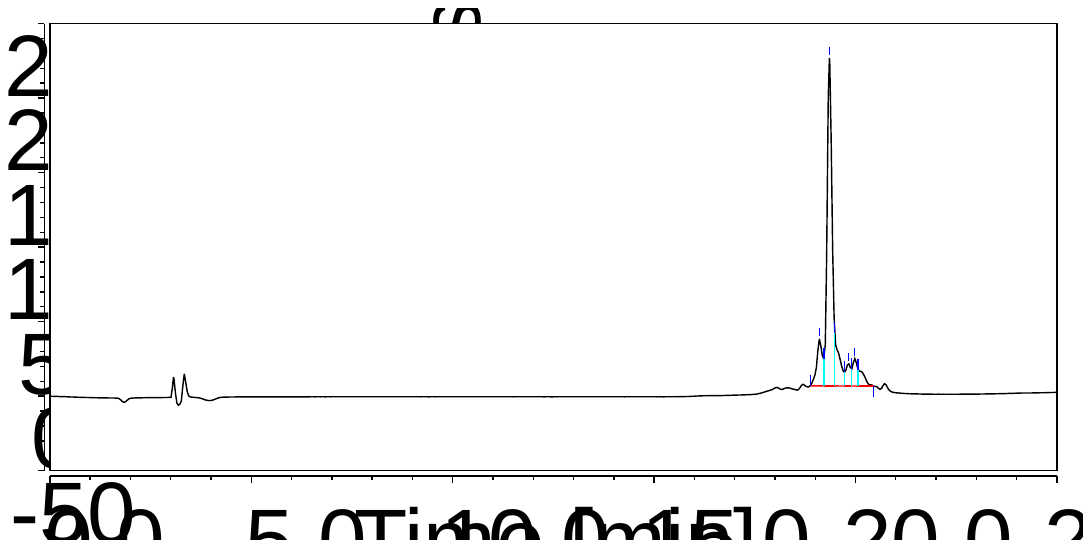

**Figure S10.** Chromatogram of fraction 10 of G. officinalis extract

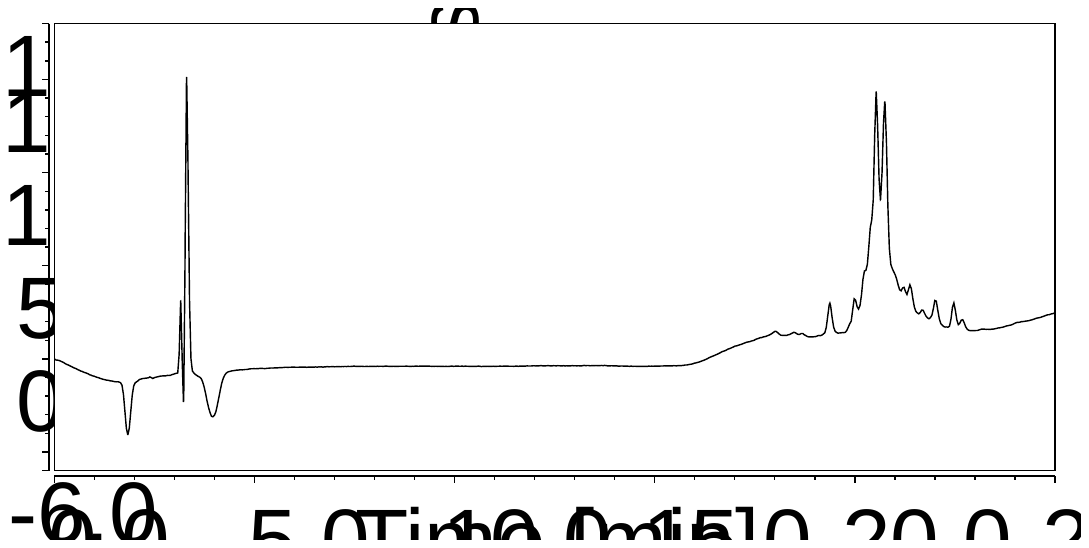

**Table S4.** Designation, content and mass of fractions of G. officinalis extract

| № previously obtained fractions | № fractions | m (extract fractions), mg |
| --- | --- | --- |
| 1 | 1 | 49.3 |
| 2, 2(2), 3, 6 | 2 | 11.8 |
| 4, 5 | 3 | 13.2 |
| 7, 7(2) | 4 | 11.8 |
| 9, 9(2) | 5 | 3.2 |

**Table S5.** Biological activity of G. officinalis extract fractions on human kidney cancer cells A498

| Indices | Control | Extraction Fractions, mg/ml | | | | | | |
| --- | --- | --- | --- | --- | --- | --- | --- | --- |
|  |  | Fraction 1  4,93  Me  (min-max)  [q25-q75] | Fraction 2  1,18  Me  (min-max)  [q25-q75] | Fraction 3  1,32  Me  (min-max)  [q25-q75] | | Fraction 4  1,18  Me  (min-max)  [q25-q75] | | Fraction 5  0,32  Me  (min-max)  [q25-q75] |
|  | After 24 hours | | | | | | | |
| Total cell count in field of vision | 123,5  (96-201) [114,5-128] | 100  (89-121)  [93-112]  p=1 | 91  (82-112)  [82-108] p=1 | 110  (92-140)  [98-121]  p=1 | | 80  (49-108)  [77-89] p=0,1549 | | 77,5  (60-94)  [73,5-87] **p=0,0477** |
| Number of dead cells in field of view | 4,5 (0-9)  [3-6] | 7 (2-12) [3-8]  p=1 | 11(6-17)  [6-16]  p=1 | 3 (0-6)  [1-3]  p=1 | | 18 (7-40)  [9-30]  **p=0,4454** | | 44,5 (31-59)  [38,5-48,5]  **p=0** |
| Ratio of dead cell count to total cell count | 0,03  (0-0,07)  [0,02-0,05] | 0,07 (0,02-0,1)  [0,03-0,09]  p=1 | 0,11 (0,07-0,2)  [0,07-0,17]  p=1 | 0,02 (0-0,06)  [0,01-0,03]  p=1 | | 0,25 (0,1-0,51)  [0,11-0,37]  p=0,4532 | | 0,58 (0,37-0,98) [0,42-0,64]  **p=0,0003** |
| Number of living cells in field of view | 118,5  (94-196)  [109-124,5] | 93 (84-118)  [85-106]  p=1 | 80 (66-98)  [73-97]  p=1 | 105  (91-138)  [95-120]  p=1 | | 60,5  (33-88)  [47-71]  p=0,1154 | | 32,5  (1-59)  [26,5-49,5]  **p=0,0009** |
| Ratio of living cell count to total cell count | 0,97  (0,93-1)  [0,95-0,98] | 0,93  (0,9-0,98)  [0,91-0,97]  p=1 | 0,89  (0,8-0,93)  [0,83-0,93]  p=1 | 0,98  (0,94-1)  [0,97-0,99]  p=1 | | 0,75  (0,49-0,9)  [0,63-0,89]  p=0,4532 | | 0,42  (0,02-0,63)  [0,36-0,58]  **p=0,0003** |
| Number of dividing cells in field of view | 2 (0-5)  [1-3] | 2 (1-3)  [1-3]  p=1 | 7 (7-13)  [7-12]  **p=0,0001** | 3 (2-6)  [2-4]  p=1 | | 7 (2-12)  [4-9]  **p=0,0032** | | 4 (2-5)  [3-4,5]  p=1 |
| Ratio of dividing cells to living cells | 0,02  (0-0,04)  [0,01-0,02] | 0,02  (0,01-0,04)  [0,01-0,03]  p=1 | 0,09  (0,07-0,16)  [0,07-0,15]  **p=0,0054** | 0,03  (0,02-0,05)  [0,02-0,04]  p=1 | | 0,11  (0,03-0,23)  [0,1-0,14]  **p=0,001** | | 0,11  (0,06-0,25)  [0,09-0,14]  **p=0,0004** |
| After 48 hours | | | | | | | | |
| Total cell count in field of view | 144  (124-174)  [130-162] | 142,5  (92-162)  [103,5-150,5]  p=1 | 37  (28-65) [31-47]  **p=0** | 138  (116-160)  [128-156]  p=1 | 58  (45-83)  [48-75,5]  **p=0** | | 35  (23-70)  [34-39]  **p=0** | |
| Number of dead cells in field of view | 24  (2-91)  [5-70] | 9 (1-23)  [4,5-15,5]  p=1 | 24 (3-37)  [16-33]  p=1 | 3 (0-14)  [0-4]  p=0,171 | 44,5 (19-52)  [35-49]  p=1 | | 26,5 (15-69)  [20-32]  p=1 | |
| Ratio of dead cell count to total cell count | 0,17  (0,01-0,58)  [0,03-0,43] | 0,07  (0,01-0,16)  [0,04-0,14]  p=1 | 0,69  (0,05-0,97)  [0,4-0,86]  p=1 | 0,02  (0-0,12)  [0-0,03]  p=1 | 0,75  (0,29-0,96)  [0,54-0,89]  p=0,4349 | | 0,69  (0,44-0,99)  [0,59-0,79]  p=0,5523 | |
| Number of living cells in field of view | 110  (54-166)  [86-142] | 124  (78-161)  [99-140]  p=1 | 17  (1-62)  [4-21]  **p=0,0042** | 138  (102-156)  [126-156]  p=1 | 15  (2-57)  [6-34,5]  **p=0,0027** | | 11  (1-19)  [10-12]  **p=0,0015** | |
| Ratio of living cell count to total cell count | 0,83  (0,42-0,99)  [0,57-0,97] | 0,93  (0,84-0,99)  [0,86-0,96]  p=1 | 0,31  (0,03-0,95)  [0,14-0,6]  p=1 | 0,98  (0,88-1)  [0,97-1]  p=1 | 0,25  (0,04-0,71)  [0,11-0,46]  p=0,4349 | | 0,31  (0,01-0,56)  [0,21-0,41]  p=0,5523 | |
| Number of dividing cells in field of view | 3  (2-7)  [3-5] | 7 (1-11)  [4,5-8,5]  p=1 | 3 (0-6)  [3-4]  p=1 | 6 (4-8)  [5-7]  p=1 | 4 (2-8)  [3-5,5]  p=1 | | 2,5 (1-5)  [1-3]  p=1 | |
| Ratio of dividing cells to living cells | 0,03  (0,02-0,06)  [0,02-0,05] | 0,05  (0,01-0,07)  [0,04-0,07]  p=1 | 0,18  (0-0,38)  [0,05-0,22]  p=1 | 0,05  (0,03-0,06)  [0,03-0,06]  p=1 | 0,23  (0,08-1,5)  [0,11-0,56]  **p=0,0024** | | 0,29 (0,07-1)  [0,25-0,3]  **p=0,0047** | |

Note: valid differences from control are shown in **bold**.

**Figure S11.** Total cell count in field of vision 24 hours after treatment with G. officinalis extract fractions

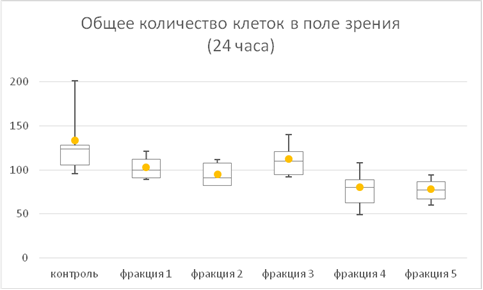

Fraction 5

Fraction 4

Fraction 3

Fraction 2

Fraction 1

Control

**Figure S12.** Ratio of dead cells to total cell count 24 hours after treatment with G. officinalis extract fractions

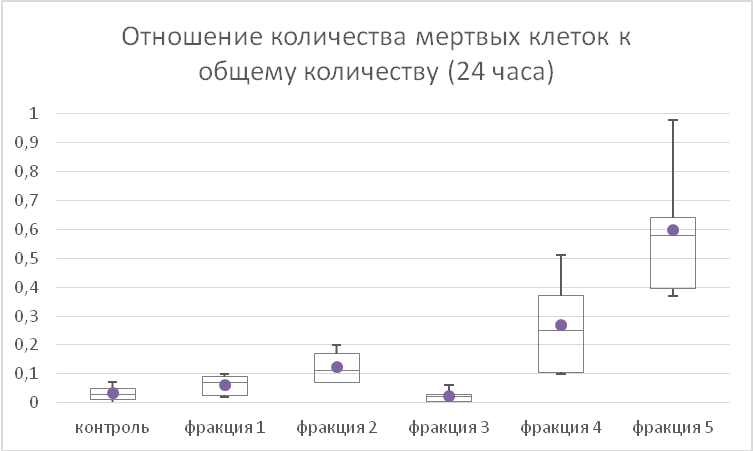

Fraction 5

Fraction 4

Fraction 3

Fraction 2

Fraction 1

Control
